# WD 40 domain of RqkA regulates its kinase activity and role in extraordinary radioresistance in *Deinococcus radiodurans*

**DOI:** 10.1101/2020.04.06.028845

**Authors:** Dhirendra Kumar Sharma, Subhash C. Bihani, Hari S Misra, Yogendra S. Rajpurohit

## Abstract

RqkA, a DNA damage responsive Serine / Threonine kinase is characterized for its role in DNA repair and cell division in *D. radiodurans*. It has a unique combination of a kinase domain at N-terminus and a WD40 type domain at C-terminus joined through a linker. WD40 domain is comprised of eight β propeller repeats held together via “tryptophan-docking motifs” and forming a typical ‘velcro’ closure structure. RqkA mutants lacking the WD40 region (hereafter referred to as WD mutant) could not complement RqkA loss in γ radiation resistance in *D. radiodurans* and lacked γ radiation mediated activation of kinase activity *in vivo*. WD mutants failed to phosphorylate its cognate substrate (e.g. DrRecA) in surrogate *E. coli* cells. Further, unlike wild type enzyme, the kinase activity of its WD40 mutants was not stimulated by Pyrroloquinoline quinine (PQQ) indicating the role of the WD motifs in PQQ interaction and stimulation of its kinase activity. Together, results highlighted the importance of the WD40 domain in the regulation of RqkA kinase signaling functions *in vivo* and thus the role of WD40 domain in the regulation of any STPK is the first time demonstrated in bacteria.

**Importance:** This study highlights the importance of the WD40 domain in activity regulation and signaling activity of bacterial serine/ threonine kinase for the first time in the bacterial response to gamma radiation and DNA damage.

## Introduction

Reversible protein phosphorylation plays an important role in transmitting extracellular signals to molecular levels by affecting numerous macromolecular events in the cell (Kobir *et al.*, 2011). The homeostasis of this process is regulated by the regulated action of protein kinases and phosphatases (Kennelly *et al.*, 1996). The most common types of phosphorylation found in proteins are on the side chains of serine/threonine, tyrosine, histidine, and aspartate residues (Hanks & Hunter, 1995). The molecular events involved in protein phosphorylation and dephosphorylation in response to DNA damage are better understood in eukaryotes as compared to prokaryotes. In bacteria, two-component system (TCS) mediated by histidine kinase and cognate response regulator is a best characterized mechanism of protein phosphorylation mediated signal transduction processes to molecular levels (Parkinson, 1993, Dutta *et al.*, 1999, Gao & Stock, 2009, Desai *et al.*, 2011, Wang *et al.*, 2008). Apart from TCS, the other mechanism of protein phosphorylation /dephosphorylation involves hank type serine threonine/tyrosine protein kinases (ST/YPKs), which phosphorylate proteins at serine, threonine and tyrosine residues (Hanks *et al.*, 1988). These ST/YPKs are known to regulate several key processes in bacterial physiology viz; abiotic stress adaptation (Pereira *et al.*, 2011, Molle & Kremer, 2010), cell morphogenesis (Macek *et al.*, 2007) and cell cycle regulation in response to DNA damage (Garcia-Garcia *et al.*, 2016, Rajpurohit & Misra, 2010). Typically, the catalytic core of such STPK folds into the two-lobed structure and the catalytic active site located in a deep cleft formed between two lobes. This type of structural conservation of the catalytic domain is found in these kinases characterized across the biological systems (Hanks *et al.*, 1988). Proteins kinases behave like a molecular switch that exists in either an “off,” inactive state or an “on,” active state (Huse & Kuriyan, 2002). The transition between “on” and “off” state is controlled by diverse mechanisms including the binding of allosteric effectors and the sub-cellular localization. For the binding of allosteric effectors, STPKs catalytic domain joined with additional domains and these extra domains. Mostly, these extra domains regulate enzyme activity through ligand-protein and protein-protein interaction. For example, PASTA repeats that associate with PrkC of *B. subtilis* that involves in spore germination (Shah *et al.*, 2008) and PknB of *M. tuberculosis* and *S. aureus* regulate cell wall morphology (Barthe *et al.*, 2010). Another type of extra domain is the C-terminal tetratricopeptide repeat domain (TPRD) found in PknG of *M. Tuberculosis* and involves in dimerization of this enzyme (Scherr *et al.*, 2007). Although, the rigid beta-propeller domains with central pore and the varying number of blades have been identified in some STPKs (Good *et al.*, 2004, Hu *et al.*, 2017), the ligand that would interact with these beta-propellers and its functional significance in the kinase activity regulation has not been studied in details.

*Deinococcus radiodurans* R1 (DEIRA) manage to survive the very high doses of DNA damaging agents including radiations with a negligible loss to its survival (Minton, 1994, Cox & Battista, 2005, Misra *et al.*, 2012). An efficient DNA double strand break (DSB) repair (Zahradka *et al.*, 2006) and a strong oxidative stress tolerance mechanism (Slade *et al.*, 2009, Blasius *et al.*, 2008, Bihani *et al.*, 2018) have been implicated to the extreme phenotypes of this bacterium. In response to DNA damage; SOS mediated DNA damage repair and cell cycle regulation is a key survival mechanism for many bacteria (Shimoni *et al.*, 2009, Bolsunovsky *et al.*, 2016). Surprisingly, *D. radiodurans* is not benefitted from LexA/RecA mediated canonical SOS response (Bonacossa de Almeida *et al.*, 2002, Narumi *et al.*, 2001). Moreover, *D. radiodurans* cells can adjust its cellular response to DNA damage by gene expression change (Liu *et al.*, 2003, Tanaka *et al.*, 2004, Rajpurohit *et al.*, 2013c) and by employing novel transcription regulator IrrE (Earl *et al.*, 2002) / by novel molecular switch PprI (Hua *et al.*, 2003), by regulating the molecular interaction of cell division and genome segregation proteins (Misra *et al.*, 2018) and by adjusting its protein homeostasis (Joshi *et al.*, 2004). Earlier, we have shown that a radiation responsive Serine /Threoninekinase (RqkA) plays a key role in DNA damage response, DSB repair and cell cycle regulation in *D. radiodurans* (Rajpurohit & Misra, 2010, Rajpurohit & Misra, 2013a, Rajpurohit *et al.*, 2016, Maurya *et al.*, 2018, Sharma *et al.*, 2020). *D. radiodurans* cells devoid of *rqkA* become hypersensitive to γ radiation and lose DSB repair ability. The domain architecture of RqkA showed an STPK domain at N-terminal and an array of β propeller motifs in WD40 domain at C-terminal (hereafter referred to as WD40 domain), and both are held together by the flexible Juxta linker region (hereafter referred to as JLR). The role of kinase domain of RqkA in γ radiation resistance of *D. radiodurans* has been demonstrated earlier (Rajpurohit & Misra, 2010, Rajpurohit & Misra, 2013a). However, the role of WD40 domain in RqkA function has not been studied yet and would be worth investigating. Here, we report the involvement of WD40 domain in the regulation of RqkA kinase function in response to γ radiation and the phosphorylation of its cognate substrate DrRecA. We demonstrated that WD40 domain deletion mutants of RqkA are severely compromised in RqkA functions in γ radiation survival and its activity stimulation by PQQ and gamma radiation. These results together suggested that PQQ interacts through WD40 domain, which seems to be responsible for both PQQ and gamma radiation response of RqkA in this bacterium and highlights WD40 domain role in the regulation of RqkA functions in *D. radiodurans.*

## Results

### 1. C-terminal domain of RqkA has structural properties of typical WD40 domain

Raptor-X server was used to prepare the homology model of RqkA. The homology model of RqkA kinase showed the presence of two distinct domains connected through a long, flexible linker region (Fig. 1A). While the N-terminal domain (Residues 1-280) defines the STPK domain, the C-terminal domain (residues 305-668) folds into eight-bladed β-propeller structure belonging to the WD40 protein subfamily. Both domains joined by flexible JLR region (Residues from 281-304). The N-terminal kinase domain shares all the structural features including the ATP binding site and other conserved motifs like P-loop, Helix-C, DFG motif, and catalytic loop as known in other Hank Type kinases (Hanks *et al.*, 1988). Important residues in the ATP binding site e.g. K42, N142, M154, D155 and the activation loop are conserved (Fig S1) “(for reviewers information only)”. Among these sites, K42 was tested as a kinase minus mutant of RqkA experimentally (Rajpurohit & Misra, 2013a). The C-terminal domain of RqkA folds into a super barrel structure consisting of eight four-stranded anti-parallel β-sheets arranged radially around a pseudo-eight-fold symmetry axis (Fig 1B). This structural arrangement is commonly known as β-propeller with four-stranded (A to D) anti-parallel β-sheets representing a blade of the propeller. By convention, each β-strand in a blade is labeled A through D with the A strand being closest to the pseudo-symmetry axis. Like many other WD40 proteins, RqkA WD40 domain creates a “velcro” closure of the ring by joining the first β-strand (strand D shown in blue in Fig. 1B) with the three strands from the last β-sheet *i.e.* blade no. 8 at C-terminus (A–C, shown in red in Fig.1B). Similar to other propellers, RqkA kinase propeller blades are connected through longer loops called “DA” loops. Among them, the DA loop connecting strand D5 to A6 (shown in yellow in Fig. S2) “(for reviewers information only)” is relatively longer than other loops and protrudes out toward the top surface. The D strands also contain charged residue in the middle and a β-bulge created by the Trp-docking motif (Fig. 2C). The tryptophan residues present at the beginning of the D strand of blade form hydrogen-bonding interactions with the main chain of two neighboring blades and form a stabilizing girdle (Fig. S3) “(for reviewers information only)”. This type of arrangement of β-propeller maintain WD40 domain structural rigidity and provide large surface for protein-protein interaction (Albrecht & Zeth, 2011). Together, RqkA model featured it as a regulatory kinase and its WD40 domain with flexible JLR region may impart in its activity regulation.

**Figure 1:**
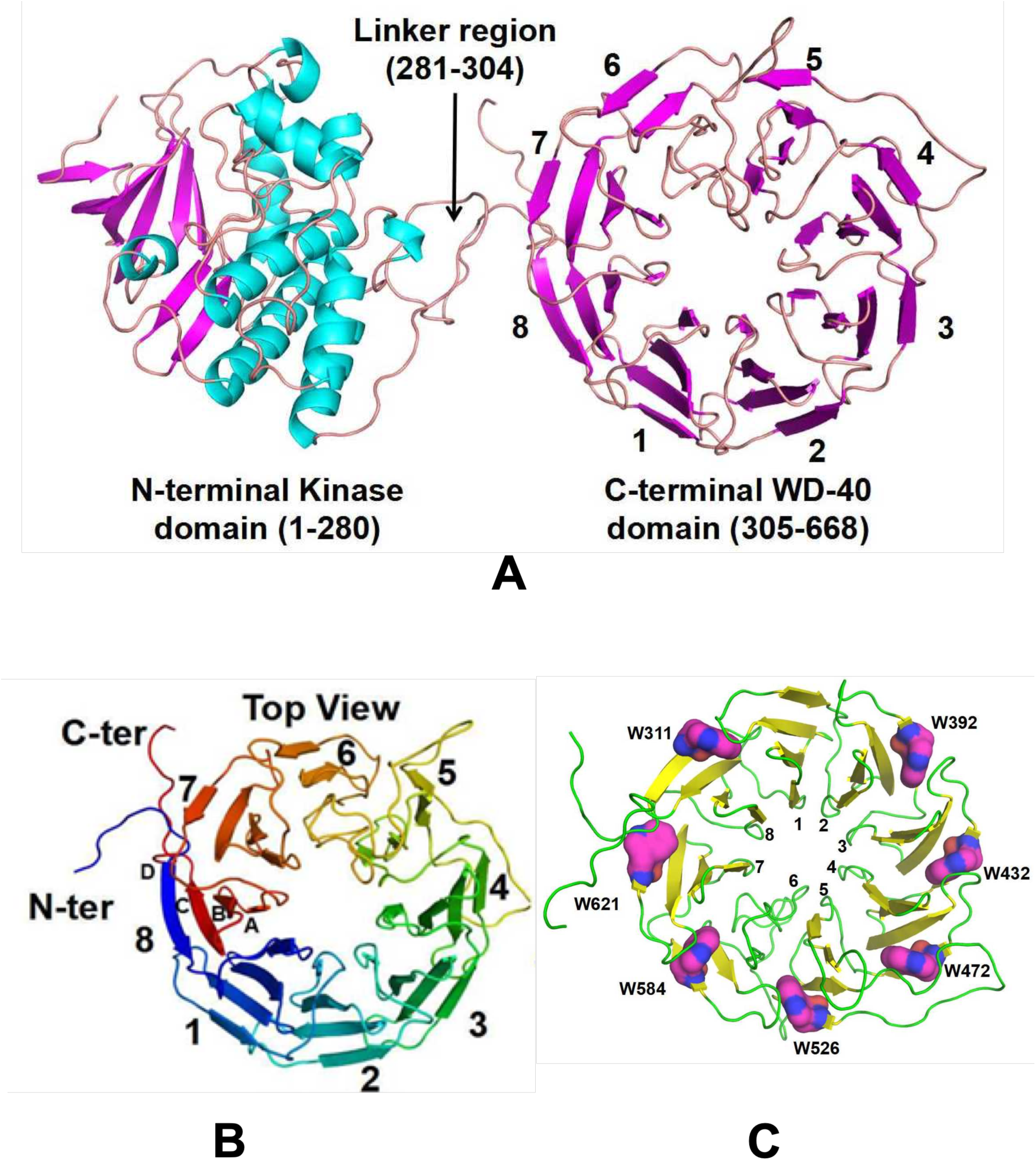
Structural features of RqkA. (A) 3-dimensional structure model of RqkA showing N-terminal kinase domain and C-terminal WD 40 domain connected with a linker. (B) Top views of RqkA WD-40 domain showing 8 bladed β-propeller with velcro closure. (C) Conserved tryptophan in 7 out of 8 blades forms stabilizing girdle.

**Figure 2:**
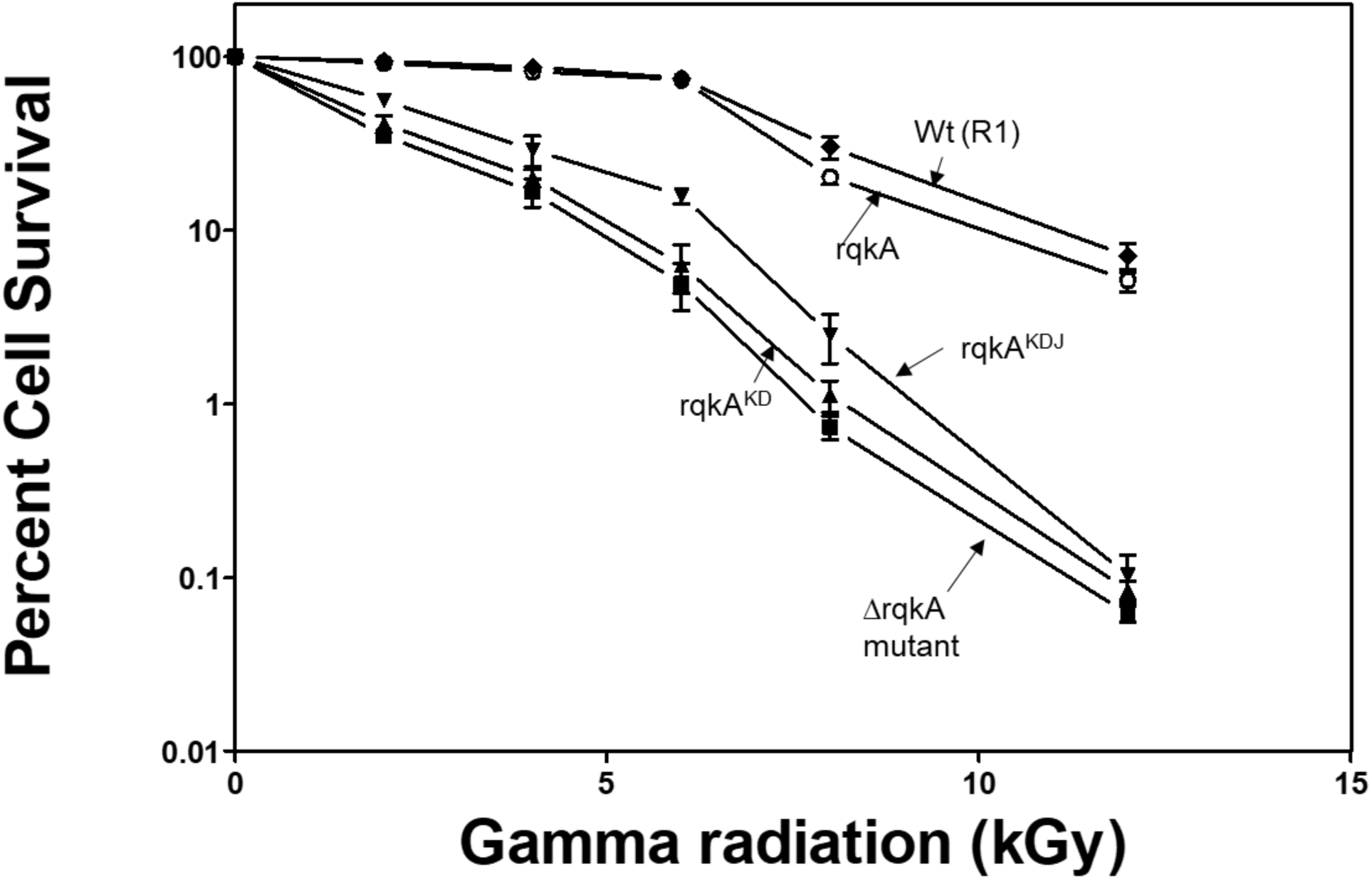
Functional complementation of RqkA loss in *ΔrqkA* mutant of *D. radiodurans* by wild type and WD40 domain mutants of RqkA. Wild type RqkA and its mutants like RqkA^KD^ and RqkA^KDJ^ were expressed in *ΔrqkA* mutant of *D. radiodurans* and the cell survival was monitored at different doses of γ radiation and compared with wild type (R1). Data given are representatives of the reproducible experiments repeated 3 times independently.

### 2. WD40 domain of RqkA kinase is required for its role in γ radiation resistance in *D. radiodurans*

Earlier, it was shown that the *rqkA* mutant of *D. radiodurans* became hypersensitive to gamma radiation and could not reassemble the shattered genome during its post-irradiation recovery (Rajpurohit & Misra, 2010). The K42A kinase mutant of RqkA had failed to complement RqkA loss in radioresistance of this bacterium (Rajpurohit & Misra, 2013a). The role of the WD40 domain in the regulation of the RqkA kinase function was further studied. Two variants of RqkA with only RqkA kinase domain ranging from 1-280 amino acids (hereafter referred to as RqkA^KD^) and RqkA kinase domain with JLR region from 1-305 amino acids (hereafter referred as RqkA^KDJ^) were generated. These were expressed *in trans* in *rqkA* deletion mutant of *D. radiodurans* and functional complementation was compared with wild type RqkA (Fig. 2). The RqkA^Wt^ complemented nearly complete to the loss of RqkA in γ radiation resistance. However, both RqkA^KDJ^ and RqkA^KD^ variants that lacked the WD40 domain but possess kinase domain with and without JLR linker respectively did not complement fully to RqkA loss of γ radiation resistance. Surprisingly, the Δ*rqkA* cells expressing RqkA^KD^ offers better support to γ radiation resistance than cells expressing RqkA^KDJ^ (Fig. 2). These results signify the role of the WD40 domain in RqkA’s *in vivo* functions. Curiously, the functional interaction of the separated domain of RqkA was also checked by co-expressing kinase and WD40 domain on the plasmid. These cells could not restore RqkA loss of radioresistance (data not given). Thus, the functional complementation studies suggested an important role of the WD40 domain of RqkA kinase in γ radiation resistance of *D. radiodurans* and highlighted the importance of kinase domain and WD40 domain being together for RqkA function.

### 3. PQQ is required for gamma radiation stimulation of RqkA phosphorylation

RqkA is characterized as DNA damage and radiation responsive kinase with its ability to sense DNA damage resulting in activation of its autokinase activity (Rajpurohit & Misra, 2010). The WD40 domain of RqkA has a similarity with the BamB protein of *E. coli* and methoxatin dehydrogenase of *Methanococcus* (Albrecht & Zeth, 2011, Anthony *et al.*, 1994). Methoxatin dehydrogenase of *Methanococcus* and ethanol dehydrogenase of *Pseudomonas* interact with PQQ through their WD40 domain (Anthony *et al.*, 1994, Schrover *et al.*, 1993). The *D. radiodurans* cells synthesize PQQ and the mutant devoid of PQQ become sensitive to radioresistance and the DSB repair was arrested (Rajpurohit *et al.*, 2008). Here, we reasoned that PQQ, a known ligand of β propeller motifs in the WD40 domain might interact with it and could serve as a regulator of kinase function in RqkA. To test this hypothesis, the *in vivo* phosphorylation of *in trans* expressed wild type RqkA and its variants RqkA^KD^ and KqkA^KDJ^ were checked in 6kGy irradiated cells and compared with unirradiated SHAM controls. Total cell-free extracts of these cells were immunoprecipitated using RqkA antibodies and the phosphorylation status of immunoprecipitate was detected using phosphor-threonine epitope antibodies. Results showed that RqkA is phosphorylated in unirradiated cells (Fig. 3, Panel (A), lane UI), which increased further upon irradiation, and remained high till 3 h of post-irradiation recovery (PIR) period and then gradually decreases to background levels (Fig.3, (A), lane 1,3,5 PIR, *ΔrqkA*). Interestingly, RqkA expressed on plasmid showed phosphorylation in *ΔpqqE* cells lacking PQQ under normal conditions. However, there was no stimulation of RqkA phosphorylation upon gamma radiation exposure and the typical kinetics of RqkA phosphorylation as seen in wild type cells during PIR was not observed in cells devoid of PQQ (Fig. 3, (A), lane 1,3,5 PIR, *ΔrqkAΔpqqE*). Stimulation of RqkA phosphorylation in response to γ radiation in *D. radiodurans* cells, while its absence when PQQ is not present (in *ΔrqkAΔpqqE)* would suggest that RqkA requires PQQ for its activity stimulation or in other words for signaling function in response to γ radiation.

**Figure 3:**
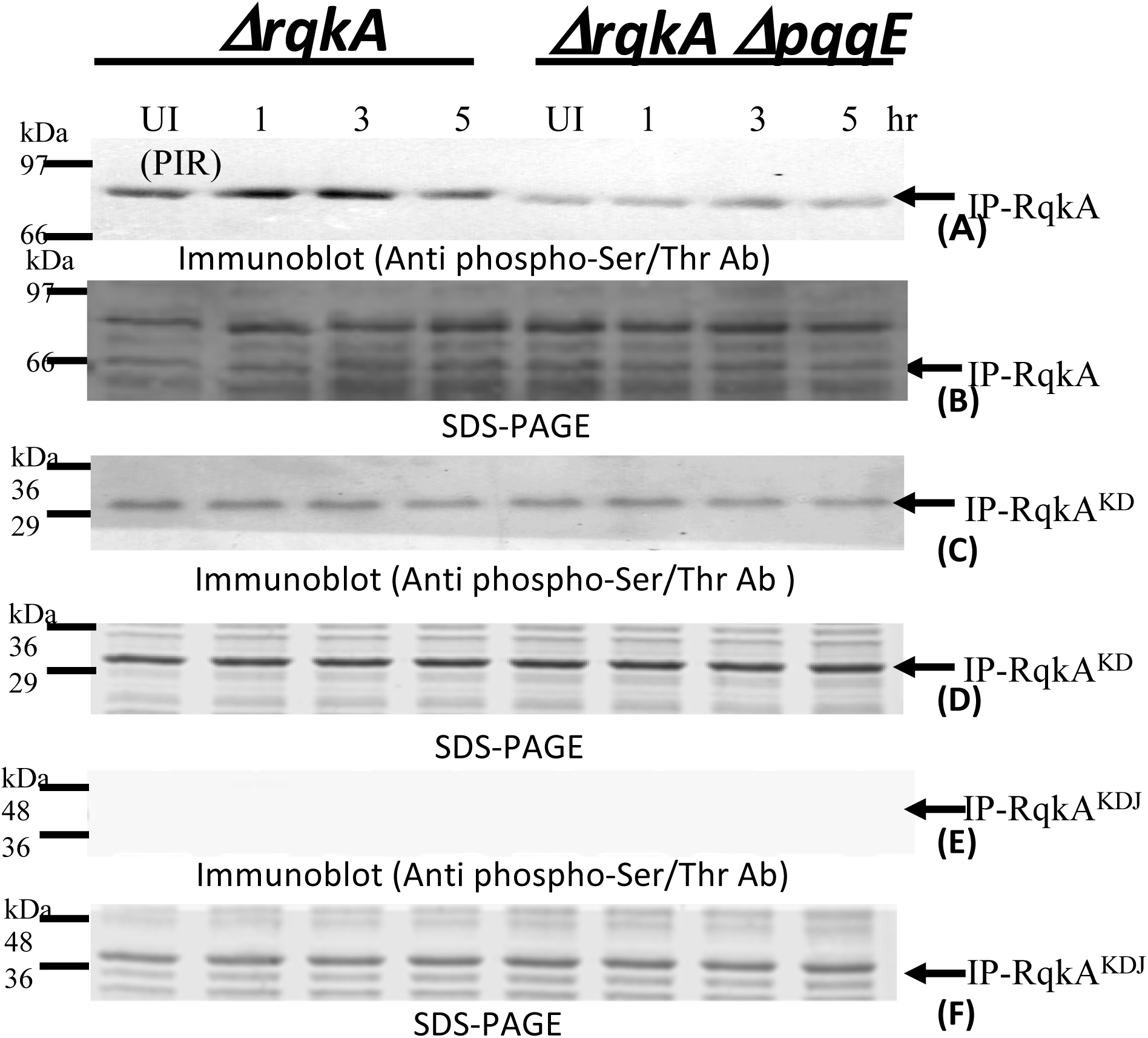
*In vivo* autophosphorylation status of RqkA and its RqkA^KD^ and RqkA^KDJ^ mutants in *Deinococcus radiodurans*. Cells were grown to exponential phase and irradiated to 6.0 kGy radiation. Gamma irradiated (I) and respective unirradiated (SHAM) control (UI) cells were grown in fresh media and aliquots were collected at different time points of post-irradiation recovery (PIR). The RqkA was immunoprecipitated from cell lysate of *ΔrqkA* and *ΔrqkAΔpqqE* mutants cells expressing either wild type RqkA or its C-terminal WD40 mutant (RqkA^KD^ or RqkA^KDJ^) using antibodies against RqkA (Anti-RqkA). Immunoprecipitants from different samples was separated on SDS-PAGE and immunoblotted using antibodies to recognize phosphor serine/threonine epitope (Anti phospho-Ser/Thr Ab) as detailed in methods. (A) *In vivo* phosphorylation status of wild type RqkA during PIR, (C) *In vivo* phosphorylation status of wild type RqkA^KD^during PIR, (C) *In vivo* phosphorylation status of wild type RqkA^KDJ^during PIR. (B), (D) and (F) showing the SDS PAGE profile of immunoprecipitated proteins using Anti-RqkA, used to probe the phosphorylation status by Anti phospho-Ser/Thr Ab of (A), (C) and (E) panel respectively. Data given are representatives of the reproducible experiments repeated 3 times.

### 4. WD40 domain regulates radiation responsiveness in the RqkA kinase function

Since PQQ interaction with RqkA was found to be through β propeller motifs in the WD40 domain, the role of the WD40 domain in gamma radiation responsiveness of RqkA phosphorylation was hypothesized and examined. For that, the WD40 deletion mutants of RqkA i.e. RqkA^KD^ and RqkA^KDJ^ were checked for phosphorylation under normal and gamma stressed conditions. The RqkA^KD^ variant showed weak phosphorylation signal under normal growth conditions but did not show γ radiation stimulation of RqkA phosphorylation under either *ΔrqkA or ΔrqkAΔpqqE* genetic backgrounds (Fig. 3, (C), lane 1,3,5 PIR). Quite interestingly, it was observed that RqkA^KDJ^ did not show phosphorylation under both normal and gamma stressed growth conditions (Fig. 3, E). The absence of kinase function in RqkA^KDJ^ while residual activity in RqkA^KD^ but both lack the γ radiation responsiveness supports the functional complementation results of γ radiation resistance. Together, these results suggest the contribution of the WD40 domain in the regulation of γ radiation responsiveness of RqkA phosphorylation and functions *in vivo.*

### 5. WD40 domain of RqkA is required for substrate phosphorylation

The mechanism that has been attributed to RqkA role in γ radiation resistance and DSB repair of *D. radiodurans* is found to be the phosphorylation of DNA repair and cell division proteins including RecA and FtsZ, and differential regulation of their functions (Rajpurohit & Misra, 2013a, Maurya *et al.*, 2018, Rajpurohit *et al.*, 2016, Sharma *et al.*, 2020). RecA is known to play an essential role in extraordinary radioresistance of *D. radiodurans* (Daly & Minton, 1996, Kim & Cox, 2002). We checked the WD40 domain role in RqkA phosphorylation of DrRecA in the surrogate *E. coli* host. RqkA and its WD40 mutants; RqkA^KD^ and RqkA^KDJ^ showed interesting phosphorylation patterns of total *E. coli* proteins and DrRecA. For instance, the RqkA expressing *E. coli* could phosphorylate endogenous proteins along with autophosphorylation of RqkA, as detected by phospho-Ser/Thr epitope antibodies (Fig. 4, K). Surprisingly, when RqkA was expressed along with its cognate substrate RecA, the majority of the phosphorylation was seen in RqkA and RecA (Fig. 4, RqkA^wt^) indicating that RqkA seems to become more specific in the presence of its cognate substrate as detected by phospho-Ser/Thr epitope antibodies (Fig. 4, compare lanes K with RqkA^wt^). However, there was no phosphorylation of RecA in cells co-expressing with either RqkA^KD^ of RqkA^KDJ^ proteins (Fig. 4, lanes RqkA^KD^ / RqkA^KDJ^). As expected, the RqkA^KDJ^ mutant did not show autophosphorylation while autophosphorylation in the RqkA^KD^ mutant has significantly reduced (Fig. 4). *E. coli* cells expressing empty *pRADgro* and *pET28a*+ plasmids showed no phosphorylation of endogenous proteins (Fig. 4, V). These results suggested that the RqkA requires its WD40 domain for trans-kinase function on its cognate substrate implying its direct or indirect involvement in enzyme-substrate interaction.

**Figure 4:**
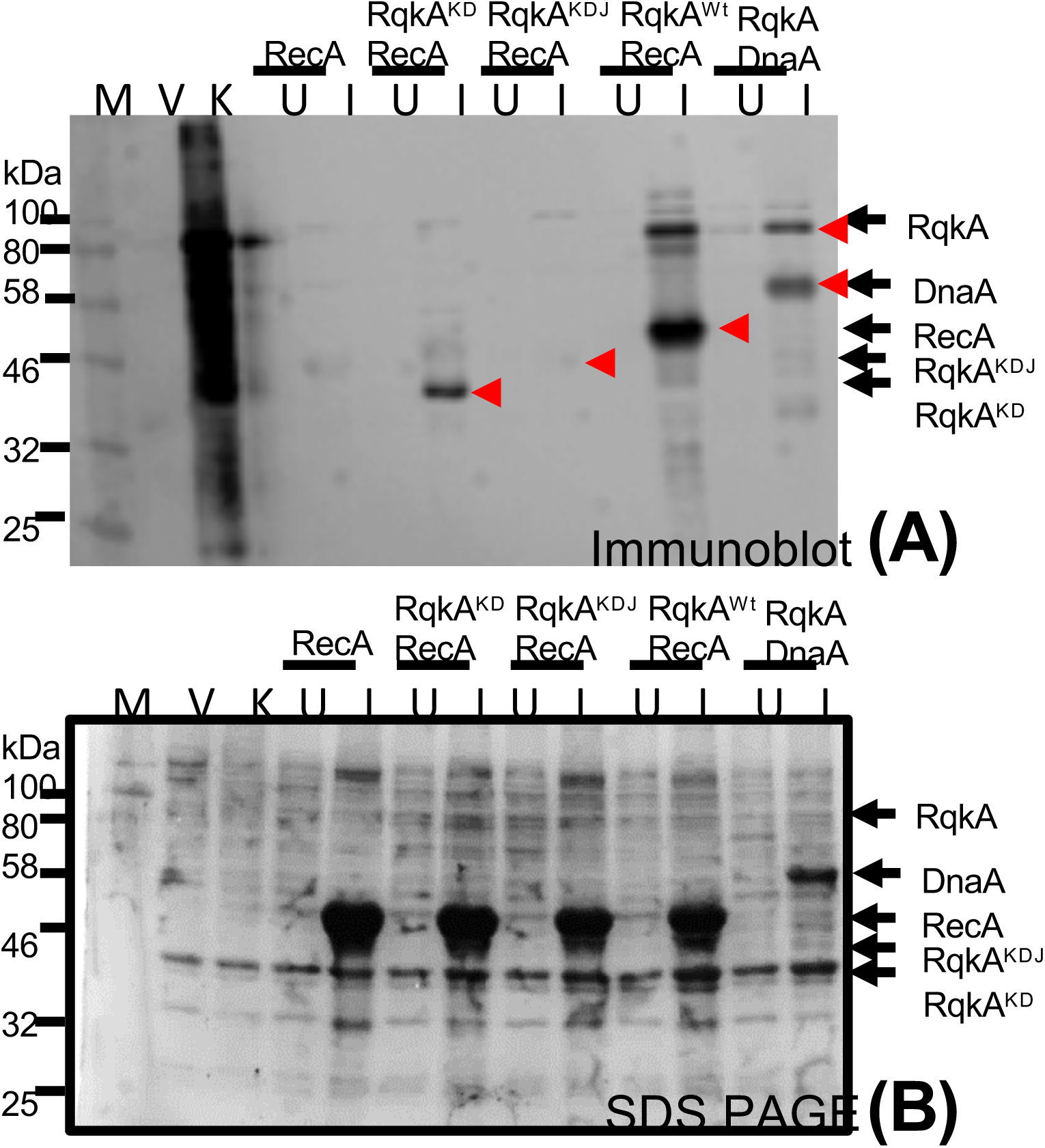
*In vivo* transphosphorylation activity of wild type RqkA and its RqkA^KD^ and RqkA^KDJ^ mutants in surrogate *E. coli* cells. For transphosphorylation studies inside the surrogate *E. coli* cells, the *E. coli* BL21 cells harboring pRAD plasmid expressing RqkA or its C-terminal mutant lacking WD40 domain (RqkA^KD^ or RqkA^KDJ^) were co-transformed with pET28a+ plasmids expressing DrRecA. pRAD vector alone used as vector control (V) while pRADrqkA expressing RqkA taken as kinase control (K). *E. coli* harboring pRADrqkA and pETdnaA plasmids expressing RqkA and DnaA were used as a positive control (Maurya et al., 2018). *E. coli* BL21 cells coexpressing kinase and its cognate substrate were grown to log phase and sampled (U), after that IPTG was added to induce the cognate substrate (DrRecA) from pET28a+ vector and sampled (I). Uninduced (U) and induced (I) cells along with vector control and kinase control cells were lysed and proteins from these cells were separated on SDS-PAGE and immunoblotted using phospho-Ser/Thr (Immunoblot (Anti p-Ser/Thr) epitopes antibodies as described in methods. Bands corresponding to phosphorylated; P-RecA, P-RqkA, P-RqkA^KD^, P-RqkA^KDJ^ and positive control (P-DnaA) are marked in the immunoblots (upper panel) (A). Sizes of immunostained protein bands were estimated using molecular weight markers (M). Arrows indicate the identity and position of respective phosphoprotein bands. Data given are representatives of the reproducible experiments repeated 3 times.

### 6. WD40 domain contributes to PQQ stimulation of RqkA kinase function

PQQ was shown to physically interact with RqkA and stimulate its kinase function *in vitro* (Rajpurohit & Misra, 2010). Many bacterial dehydrogenases have also been characterized to interact with PQQ through conserved β propeller motifs and regulate enzyme activity (Anthony & Ghosh, 1998). However, the functional implications of PQQ interaction with β propeller motifs in the WD40 domain of RqkA are not known. The RecA phosphorylation by RqkA or by its WD40 mutants RqkA^KD^ and RqkA^KDJ^ was checked in surrogate *E. coli* grown with and without PQQ (1µM). Results showed that RqkA could phosphorylate RecA, which further enhanced when PQQ was supplemented (Fig. 5, RqkA^Wt^ RecA). RqkA^KDJ^ mutant could not phosphorylate RecA irrespective of the presence of PQQ (Fig. 5, RqkA^KDJ^RecA). Notably, RqkA^KD^ mutant although showed a low level of RecA phosphorylation but did not improve in the presence PQQ (Fig. 5, RqkA^KD^RecA). These results suggested that the WD40 domain of RqkA seems to be the site for PQQ interaction and plays a decisive role in PQQ regulation of RqkA kinase activity certainly in response to gamma radiation damage.

**Figure 5:**
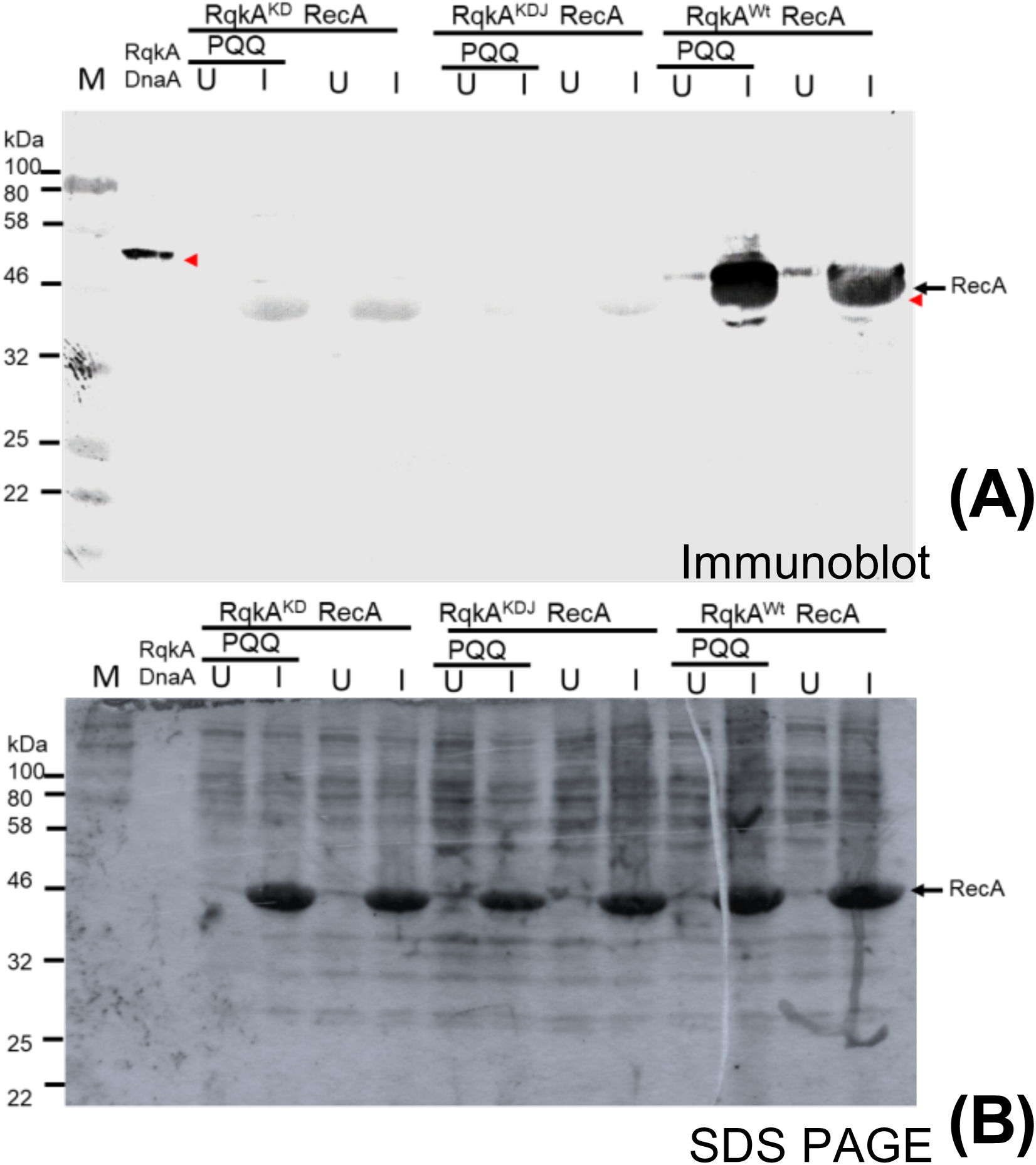
The effect of PQQ on *in vivo* transphosphorylation activity of RqkA and its RqkA^KD^ and RqkA^KDJ^ mutants in surrogate *E. coli* cells. The transphosphorylation of DrRecA by wild type RqkA kinase or by its C-terminal mutant lacking WD40 domain (RqkA^KD^ or RqkA^KDJ^) was checked in the cells supplemented with or without exogenous PQQ. *E. coli* BL21 cells coexpressing kinase and its cognate substrate (DrRecA) were grown to log phase and sampled (U) after that IPTG was added to induce the cognate substrate (DrRecA) from pET28a+ vector and sampled (I). A similar experiment was performed where cell growth supplemented with PQQ (1µM) and sampled uninduced (U) and induced (I) cells (PQQ). PQQ supplemented and no supplemented, Uninduced (U) and induced (I) cells were lysed and proteins from these cells were separated on SDS-PAGE and immunoblotted using phospho-Ser/Thr (Immunoblot (Anti p-Ser/Thr) epitopes antibodies as described in methods. Bands corresponding to phosphorylated; P-RecAand positive control (P-DnaA) are marked in the immunoblots (upper panel) (A). Sizes of immunostained protein bands were estimated using molecular weight markers (M). Arrows indicate the identity and position of respective phosphoprotein bands. Data given are representatives of the reproducible experiments repeated 3 times.

## Discussion

*Deinococcus radiodurans* is best known for its extraordinary radioresistance with a D10 dose between 10-15kGy (Misra *et al.*, 2013, Slade *et al.*, 2009). This bacterium can resist nearly 200 double-strand breaks and ∼3000 single-strand breaks per genome produced by 5000 Gy gamma radiation without loss of cell viability (Cox & Battista, 2005). The extreme phenotype of this bacterium has been attributed to its ability to protect biomolecules from oxidative damage (Daly *et al.*, 2010) and efficient DSB repair (Zahradka *et al.*, 2006, Slade *et al.*, 2009). Such a high dose of gamma radiation would produce a huge influx of oxidants causing damage to macromolecules leading to molecular responses and cell cycle regulation. Earlier, we had characterized the PQQ role in bacterial tolerance to the photodynamic effects of Rose Bengal (Khairnar *et al.*, 2003) and as an antioxidant and radioprotector *in vitro* (Misra *et al.*, 2004). PQQ role in radioresistance and DSB repair was also shown in *D. radiodurans* (Rajpurohit *et al.*, 2008). Using PQQ as a molecular link, we characterized RqkA as a cell signaling kinase having an indispensable role in cell cycle regulation in this bacterium (Maurya *et al.*, 2018, Rajpurohit *et al.*, 2016, Rajpurohit & Misra, 2010, Rajpurohit & Misra, 2013a, Sharma *et al.*, 2020). This work has gained a greater significance because of (i) the absence of LexA/RecA type canonical SOS response mechanism of bacterial DNA damage response and cell cycle regulation in this bacterium (Bonacossa de Almeida *et al.*, 2002, Narumi *et al.*, 2001), and (ii) any Ser/Thr protein kinase role in the bacterial response to DNA damage and cell cycle regulation is reported the first time.

The molecular signaling mechanism involving protein phosphorylation is ubiquitously recognized as an important regulatory process of rapid and reversible modification of the physio-chemical properties of a protein. This process triggers several possible consequences in protein biology like change in enzyme activity, oligomerization state, interaction with other proteins, subcellular localization or half-life (Kobir *et al.*, 2011). The involvement of protein kinases particularly Ser / Thr / Tyr kinase in cell cycle regulation and cellular differentiation have been extensively studied in eukaryotes (Zhou & Elledge, 2000). These signaling events ensure the effectiveness of repair enzymes for an efficient DNA strand break repair or to the demise of the cells by a regulated mechanism such as apoptosis, ensuring the genomic stability of the organism (Sancar *et al.*, 2004). Bacteria, being a simple bag of metabolites can use extensive cellular signaling networks to coordinate with cellular functions. In bacteria, the majority of environmental signals are transduced through the two-component system (TCS) involving histidine kinase and cognate response regulator, of the signal-transduction pathway (Bourret & Silversmith, 2010, Oshima *et al.*, 2002). The STPKs are widespread in bacterial genomes (Leonard *et al.*, 1998) and their role in bacterial growth (Sassetti *et al.*, 2003, Fernandez *et al.*, 2006), virulence (Didier *et al.*, 2010), persistence and reactivation (Shah *et al.*, 2008), cell division and development (Molle & Kremer, 2010, Pereira *et al.*, 2011) have been extensively documented. Many bacteria-harboring multipartite genome systems have been identified and their genome composition is heavily represented by STPKs (Krupa & Srinivasan, 2005, Leonard *et al.*, 1998, Kennelly, 2002, Misra *et al.*, 2018). However, the studies on STPK roles in the bacterial response to DNA damage, repair and cell cycle regulation has been discovered very recently. RqkA has N-terminal STPK domain and C terminal WD40 domain with an array of β propeller motifs (Fig.1A). WD40 domain is widespread in eukaryotes and has been found associated with a structurally and functionally different class of proteins ranging from signal transduction mediated by G-protein (Wall *et al.*, 1995), transcription regulation (Mylona *et al.*, 2006), ubiquitin depended protein degradation (Skaar *et al.*, 2014) and chromatin modification (Zhang & Zhang, 2015). WD proteins were suggested to be rare in prokaryotes. However, recent studies have suggested the presence of WD domain in proteins across the bacterial kingdom (Hu *et al.*, 2017). Most well-known among them is BamB and PQQ dependent alcohol dehydrogenases (Albrecht & Zeth, 2011, Anthony & Ghosh, 1998). However, there is no study linking the WD domain to any specific function. This study shows for the first time that the presence of the WD domain in RqkA is crucial for the proper functioning of its N-terminal kinase domain. When we checked the role of the WD40 domain in RqkA function in γ radiation resistance in *D. radiodurans* and the phosphorylation of its cognate substrate, the WD40 domain mutants; RqkA^KDJ^ and RqkA^KD^ failed to complement RqkA loss of γ radiation resistance as well activity stimulation in response to γ radiation (Fig. 2, 3). These results suggested the requirement of the WD40 domain in RqkA kinase function in γ radiation resistance in *D. radiodurans*. It may be highlighted that RqkA^KDJ^ and RqkA^KD^ which are different by the juxta-linker region (JLR), behaved differentially in functional complementation of RqkA loss. The JLR region of many eukaryotic receptor tyrosine kinases (c-Kit, EphB2 and Flt3) has an autoinhibitory role by blocking the substrate access to a nucleotide-binding pocket of kinase domain by phosphorylation of key tyrosine residues within JLR region (Chan *et al.*, 2003, Griffith *et al.*, 2004, Wybenga-Groot *et al.*, 2001). In RqkA also, tyrosine 287 (Y287) and threonine 281 and 294 (T281 and T294) are in the JLR region (281-305 amino acid), which raises the possibility of autophosphorylation mediated conformational change of JLR region and a possibility of attenuation of RqkA function.

STPKs are believed to be promiscuous and can phosphorylate proteins non-specifically *in vitro*. However, if this is true *in vivo*, then overexpression of STPKs should be lethal to be survival of the host, which has not been observed in most of the cases studied so far. Here, we observed that when *E. coli* expressing only RqkA, many host proteins are phosphorylated, which reduced drastically in the presence of its cognate substrate from *D. radiodurans* (Fig. 4). This could be explained on the assumption that RqkA in the presence of its substrate becomes more specific and that would be possible only when specific substrates interact with enzyme through unique contact regions in the kinase, thus the WD40 domain of RqkA may provide such specific interaction sites. The WD40 domain in RqkA showed several structural features of typical β-propeller proteins. RqkA appears to be related to both BamB and PQQ-dependent dehydrogenases (PQQ-DHs), sharing features such as the eight bladed β-propeller fold and the presence of repeating “tryptophan-docking motifs” (Fig. 1C). β-propeller blades of the WD40 domain in RqkA are joined together with DA loops providing a major molecular surface on the top surface (Fig.2B). The surface of the blades of WD proteins has been shown to take part in protein-protein interactions (Wu *et al.*, 2012). Interestingly, DA loop connecting strand D5 to A6 (DA^56^) is longer than other loops and protrudes out toward the top surface which may provide an additional surface for some specific interactions (Fig. S2) “(for reviewers information only)”. PQQ has been shown to physically interact with RqkA kinase (Rajpurohit *et al.*, 2010) and binding of PQQ with the WD40 domain of many dehydrogenases help them to attain holoenzyme activity (Anthony *et al.*, 1994, Schrover *et al.*, 1993). Our results of PQQ mediated stimulation of wild type RqkA but not of WD40 mutants of RqkA argue in favor of WD40 domain and may provide large surface for the specific interaction and help in enhance its kinase activity (Fig.5). Thus, the possibility of WD40 domain role in determining the substrate specificity of RqkA in its kinase function is suggested.

The ionizing radiation damages biological macromolecules indirectly through the radiolysis of water producing reactive oxygen and nitrogen species, and directly by the deposition of ionizing energy on covalent bonds and their breakage. The metabolic cross-talk between both these effects has been studied by measuring macromolecular damages as a function of antioxidant metabolites (AOM), antioxidant unique peptides, and antioxidant enzymes (AOE) (Smith *et al.*, 2017, Reisz *et al.*, 2014). The role of AOM in direct regulation of macromolecular events associated with DNA damage response and repair have not been studied elaborately. This study has brought forth some experimental evidence to delineate molecular links between oxidative stress and DNA damage response and repair, albeit in a restricted bacterial model system *Deinococcus radioduarns*. Earlier, PQQ has been shown to react with artificially produced reactive oxygen species and function as antioxidant and radioprotector (Misra *et al.*, 2004, Misra *et al.*, 2012). Pyrroloquinoline quinone (PQQ) identified in this bacterium and having a role in oxidative stress tolerance as well as act as an inducer to gamma radiation responsive Ser/Thr kinase (RqkA) and regulates extraordinary radioresistance and DSB repair in this bacterium (Rajpurohit *et al.*, 2008, Rajpurohit and Misra, 2010). The β propeller motif has been shown to involve in interaction with PQQ in dehydrogenases that require it as a coenzyme for their activity (Anthony *et al.*, 1994, Schrover *et al.*, 1993). Ligand binding analyses using the Raptor-X server also suggested the top surface as a potential binding site for PQQ. When the gamma radiation and DNA damage responsiveness of the WD40 mutant of RqkA was measured as a function of autophosphorylation of RqkA and its WD40 mutant, WD40 mutant failed to respond both gamma radiation and DNA damage (Fig.3, C, E). Similarly, RqkA autophosphorylation was not stimulated in the *ΔpqqE* mutant (devoid of PQQ) in *D. radiodurans* (Fig.3, A, *ΔpqqE*). The loss of γ radiation mediated activation of RqkA in the *ΔpqqE* mutant and absence of γ radiation response of RqkA^KD^ autophosphorylation suggest the importance of both WD40 domain and PQQ in the activation of RqkA function in response to γ radiation and DNA damage-induced signaling through RqkA. PQQ has recently been shown to physically interact with cellular proteins in the mammalian system and crucial for the regulation of their enzymatic activity (Akagawa *et al.*, 2016). RqkA phosphorylates DNA repair and cell division proteins in response to gamma radiation and DNA damage (Rajpurohit and Misra, 2013a, Rajpurohit *et al.*, 2016, Maurya *et al.*, 2018). Recent study has shown the presence of the WD40 domain in more than 4000 prokaryotic proteins where the majority of these proteins are STPKs (Hu *et al.*, 2017). Here, our results emphasize the role of the WD40 domain in activity regulation of STPK (RqkA) and the bacterial response to gamma radiation damage. Based on the finding of this study and our earlier studies we propose a model of RqkA kinases activation in response to γ radiation (Fig.6). It summarizes that γ radiation causes the cellular DNA damage and reactive oxygen and nitrogen species (ROS and RNS) generation, which leads to the upregulation of PQQ synthesis in *D. radiodurans* cells. PQQ activates the DNA damage responsive RqkA kinase through its interaction with conserved the WD40 domain and this interaction activates RqkA autophosphorylation mediated activation and support of radiation survival (Fig.6, A). In the absence of WD40 domain; RqkA could not be activated by PQQ leading RqkA inability to support radiation survival (Fig.6, B).

**Figure 6:**
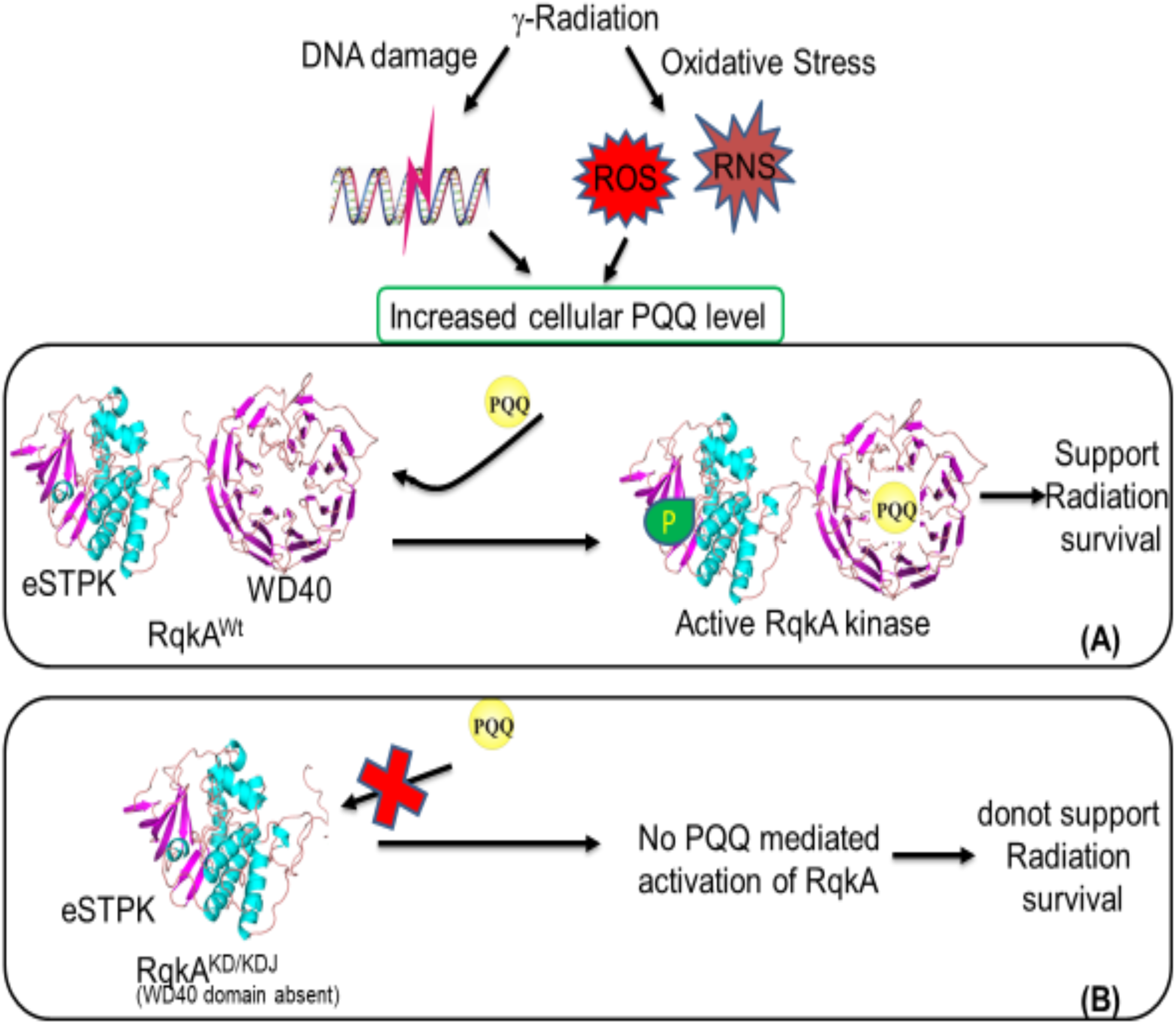
A model showing the activation of RqkA kinase. **A** DNA damage responsive kinase activation depends on its interaction with PQQ through conserved WD40 domain. The level of PQQ strongly elevated after γ radiation / or under oxidative stress which leads to binding with the C-terminal WD40 domain of RqkA kinase and this interaction activates RqkA autophosphorylation mediated activation and support of radiation survival (Panel A). In the absence of WD40 domain, RqkA could not able to activate by PQQ leads to RqkA inability to support radiation survival (Panel B).

In conclusion, we report the functional characterization of the C-terminal WD40 domain of RqkA in the regulation of the signaling function of RqkA. We demonstrated that the WD40 domain is the site for PQQ mediated activation of RqkA, which could provide a molecular link between oxidative stress response and DNA damage response mediated by PQQ and RqkA in *D. radiodurans* respectively. The inability of WD40 mutant to phosphorylate a representative substrate DrRecA of RqkA, as well as the lack of enhancement of DrRecA phosphorylation by RqkA in the absence of PQQ suggested the role of WD40 and PQQ in the regulation of RqkA. While independent studies would be required to completely delineate a direct link between WD40 domain and PQQ interaction, the stoichiometry of PQQ interaction with RqkA and its impact on *in vivo* signal transduction in response to DNA damage. The available results suggest the role of the WD40 domain and PQQ in the regulation of signaling activity of RqkA and to the best of our knowledge, this is the first report on the role of the WD40 domain in the regulation of the STPKs in the bacterial response to gamma radiation and DNA damage.

## Experimental procedures

### Bacterial strains, plasmids, and materials

*D. radiodurans* R1 (ATCC13939) was grown in TGY (Bacto tryptone (1%), Glucose (0.1%) and Yeast extract (0.5%)) medium with shaking at 180 rpm at 32°C. *E. coli* strain NOVABLUE was used for cloning and maintenance of all the plasmids; *E. coli* strain BL21 (DE3) pLysS was used for the expression of recombinant proteins. *E. coli* cells harboring pRADgro and pET28a(+)were maintained in the presence of required antibiotics. The pRADgro and their derivatives were maintained in the presence of ampicillin (100 µg/ml) in *E. coli* and chloramphenicol (8 µg/ml) in *D. radiodurans*as described previously (Misra *et al.*, 2006). Standard protocols for all recombinant techniques were used as described in (Green and Sambrook, 2012). An antibody against phosphor serine/threonine epitope was procured commercially (Cell Signaling Technology, USA). Antibodies against RqkA of *D. radiodurans* were commercially produced in the rabbit (MERCK Millipore, India). Molecular biology grade chemicals and enzymes were procured from Sigma Chemicals Company, USA, Roche Biochemicals, Mannheim, Germany, New England Biolabs, USA, and Merk India Pvt. Ltd. India.

### Homology Modelling and Analyses

Homology modelling was performed by the Raptor-X server (Källberg *et al.*, 2012). RaptorX server used multiple templates to build N and C-terminal domains. The N-terminal kinase domain was modeled using templates like PknA of *Mycobacterium tuberculosis* (PDB Ids 6B2Q, 4OW8 and 4×3F), PknB of *Mycobacterium tuberculosis* (PDB Id 3ORI) and Staphylococcus aureus (PDB Id 4EQM). Similarly, *Escherichia coli* BamB protein (PDB Ids 3P1L, 4PK1), *Pseudomonas aeruginosa* BamB protein (PDB Id 4HDJ), *Moraxella catarrhalis*BamB protein (PDB Id 4IMM) and *Methylomicrobium buryatense* methanol dehydrogenase (PDB Id 6DAM) were used to build C-terminal region. The model built by Raptor-X was then refined using the ReFold server (Shuid *et al.*, 2017). The geometry of the model was optimized automatically using Phenix and manually using WinCoot. The quality of the optimized model was then evaluated using ProSA, QMEAN and PROCHECK (Wiederstein & Sippl, 2007, Benkert *et al.*, 2011, Laskowski *et al.*, 1993). Ligand binding sites in the model were also identified using Raptor-X server. Conserved residues in the rqkA were analyzed using the Consurf server (Ashkenazy *et al.*, 2016). The surface electrostatic potentials of the structure were generated using APBS software with default settings as implemented in PyMol. For the assessment of the quality of the model, several validation software’s were used. The Raptor-X server uses many parameters to judge the quality of the model. The P-value, uGDT and GDT scores for the alignment of query with the top ranked template are used together to assess the quality of the resulting model structure. The P-value, uGDT and GDT for RqkA model were 5.57^e-15^, 391 and 58 respectively. A good P-value is <10^−3^ for mainly alpha proteins and < 10^−4^ for mainly beta proteins. For a protein with >100 residues, uGDT>50 is a good indicator. For a protein with <100 residues, GDT>50 is a good indicator. GDT is calculated as uGDT divided by the protein (or domain) length and multiplied by a 100. uGDT (GDT) measures the absolute model quality while P-value evaluates the relative quality of a model. The resulting model has good values of all three parameters suggesting the good quality of the model. The model was also analyzed for correct stereochemistry using PROCHECK. Ramachandran plot analysis using PROCHECK shows that 99.4% of the amino acid residues are in the allowed region and 0.6% are in the generously allowed region. No residue is found to be in the disallowed region. Pro-SA analysis of the model showed a Z-score of −9.74 which is in the range usually found for experimental protein structures of similar sizes. Analysis using the Verify-3d server also showed good model quality with 95.36% of the residues having averaged 3D-1D score >= 0.2.

### Construction of WD40 mutants

The Genomic DNA of *Deinococcus* was prepared as published previously (Battista *et al.* 2001). For the cloning of *rqka*^Wt^, *rqka*^KD^ and *rqka*^KDJ^ genes in pRADgro plasmid (Rajpurohit and Misra, 2013b), DNA fragments were PCR amplified from the genomic DNA of DEIRA using primers listed in Table 1. PCR product was ligated at *ApaI* and *XbaI* sites in pRADgro to yield pGrorqka, pGrorqka^KD^, and pGrorqka^KDJ^. Plasmid DNA was prepared from these clones and the presence of insert in these plasmid samples was confirmed by restriction analysis and by sequencing. The recombinant plasmid was transformed into *D. radiodurans* and chloramphenicol resistant clones were isolated on TGY agar plates containing chloramphenicol (5μg/ml).pGrorqka^Wt^, pGrorqka^KD^, and pGrorqka^KDJ^ plasmids were also transformed to *E. coli* cells expressing pETrecA (Rajpurohit *et al.*, 2016) for transphosphorylation studies in surrogate *E. coli* cells. These cells were grown in LB medium supplemented with Ampicillin (100µg/ml) and Kanamycin (25 µg/ml).

**Table 1.**
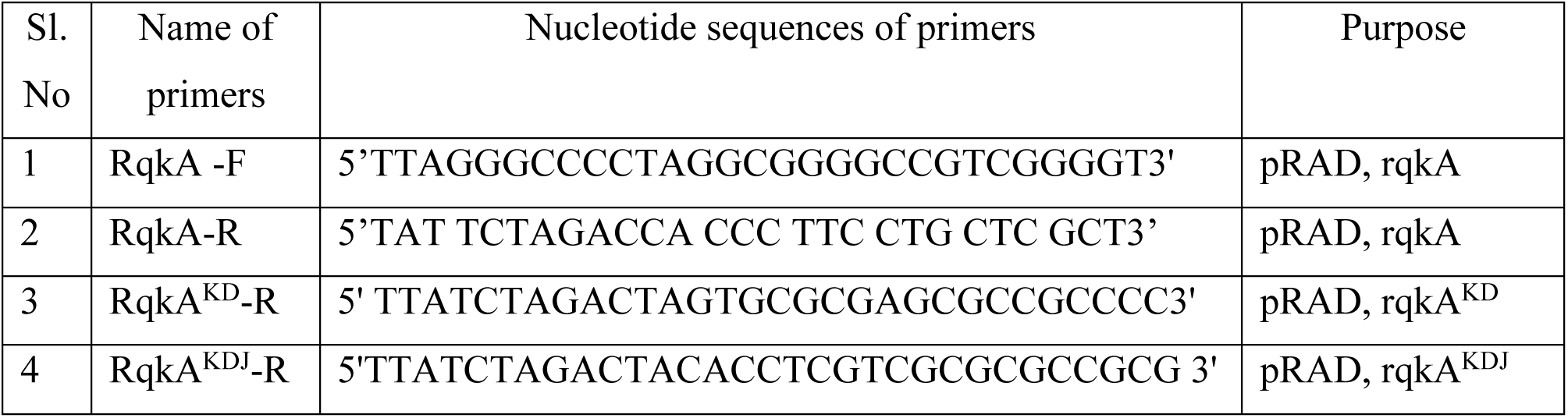
List of primers used.

### *In vivo* phophorylation studies in *D. radiodurans*

Phosphorylation studies were carried out in *D. radiodurans* as described earlier (Rajpurohit and Misra, 2010). For that, the *ΔrqkA* and *ΔrqkAΔpqqE* mutants of *D. radiodurans* expressing wild type RqkA or its WD40 mutants (RqkA^KD^ or RqkA^KDJ^) on plasmid were irradiated with 6 kGyγ radiation and allowed to recoverin TGY medium as described earlier (Mishra *et al.*, 2019). Different aliquots were collected during post-irradiation recovery (PIR) and its corresponding SHAM controls, washed with 70% ethanol and snap-frozen in liquid nitrogen before storing at −70 ° C overnight. For measuring the levels of autophosphorylation in RqkA and its WD40 mutants like RqkA^KD^ or RqkA^KDJ^, the cell-free extracts were made and immunoprecipitated using polyclonal RqkA antibodies followed by immunoblotting using phospho-Ser / Thr antibodies (catalog no. 9631S, Cell Signaling Technology, USA) as described earlier (Maurya *et al.*, 2016). In brief, the cells were treated with lysozyme (10 mg/ml) for 1 h at 37°C, followed by 0.5% NP-40 in cell lysis buffer (20 mM Tris-HCl [pH 8.0], 50 mM NaCl, 1 mM PMSF, 1 mM DTT). Treated cells were disrupted by either by sonication on an ice bath for 1 min and cleared supernatant was obtained by centrifuging at 12000g for 30 min. Approximately, 500µg total proteins in cell-free extract incubated with RqkA antibodies raised in rabbit in binding buffer (140mM NaCl, 8 mM sodium phosphate, 2mM potassium phosphate, and 10mM KCL, pH 7.4). The mixture was incubated overnight at 4° C and to this; the Protein G agarose beads were added. The content was passed through the Econopack column (Biorad, USA) and washed thrice with binding buffer and eluted with 500mM NaCl in binding buffer. The eluent was precipitated with 2.5 volume of ice-chilled acetone and precipitate was dissolved in 2X Laemmli buffer for SDS-polyacrylamide gel electrophoresis. Proteins were separated and on 10% SDS-PAGE gel and blotted onto PVDF membrane and probed with polyclonal phospho-Ser/Thr antibodies (Cell Signaling Technology, USA). Signals were detected using anti-rabbit IgG conjugated with alkaline phosphatase and the color reaction substrates NBT-BCIP (Roche Biochemicals, Germany).

### Phosphorylation studies in surrogate *E. coli*

The phosphorylation of DrRecA by RqkA or its WD40 mutants (RqkA^KD^ or RqkA^KDJ^) was checked using *E. coli* surrogate host co-expressing these proteins from plasmids as described earlier (Maurya *et al.*, 2018). For that, *E. coli* BL21 (DE3) pLysS cells were co-transformed pRADrqkA, pRADrqkA^KD^ and pRADrqkA^KDJ^ with pETrecA, separately. The recombinant proteins were induced with IPTG and an equal number of cells expressing wild type RqkA or RqkA^KD^ and RqkA^KDJ^ with DrRecA were lysed in 2X Laemmli buffer. The clear supernatant was separated on SDS-PAGE and transferred to PVDF membrane and probed with polyclonal phosphor-Ser/Thr epitope antibodies (Cell Signaling Technology, USA) as described earlier (Rajpurohit *et al.*, 2016). To see the PQQ effect on transphosphorylation activity of RqkA, the cells were grown in the presence of 1µM PQQ and compared with cells grown without PQQ for DrRecA phosphorylation as discussed above.

## Acknowledgements

Acknowledge the Department of Atomic Energy (DAE), Government of India for financial and structural support.

## Author contribution

D.K.S. planning and conducting experiments, results analysis, discussion. S.C.B. did all computational and modeling work. H.S.M. results analysis and discussion, paper writing and co-correspondence. Y.S.R. conceived the study, planning and execution of experiments, results analysis, discussion, paper writing, communication and correspondence for publication.

## Supplementary data

**Figure S1:**
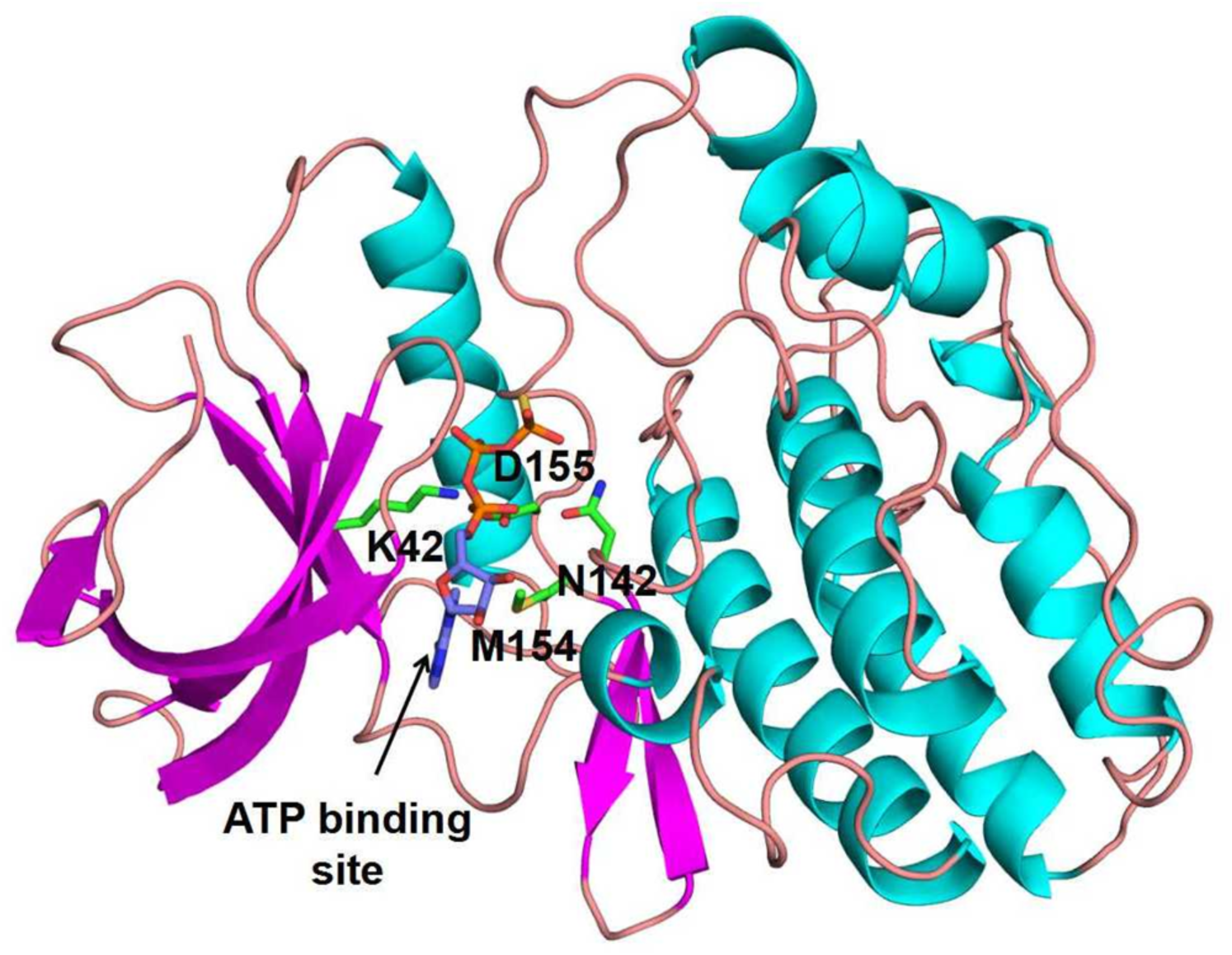
3-dimensional structural model of kinase domain of RqkA with the predicted ATP binding site.

**Figure S2:**
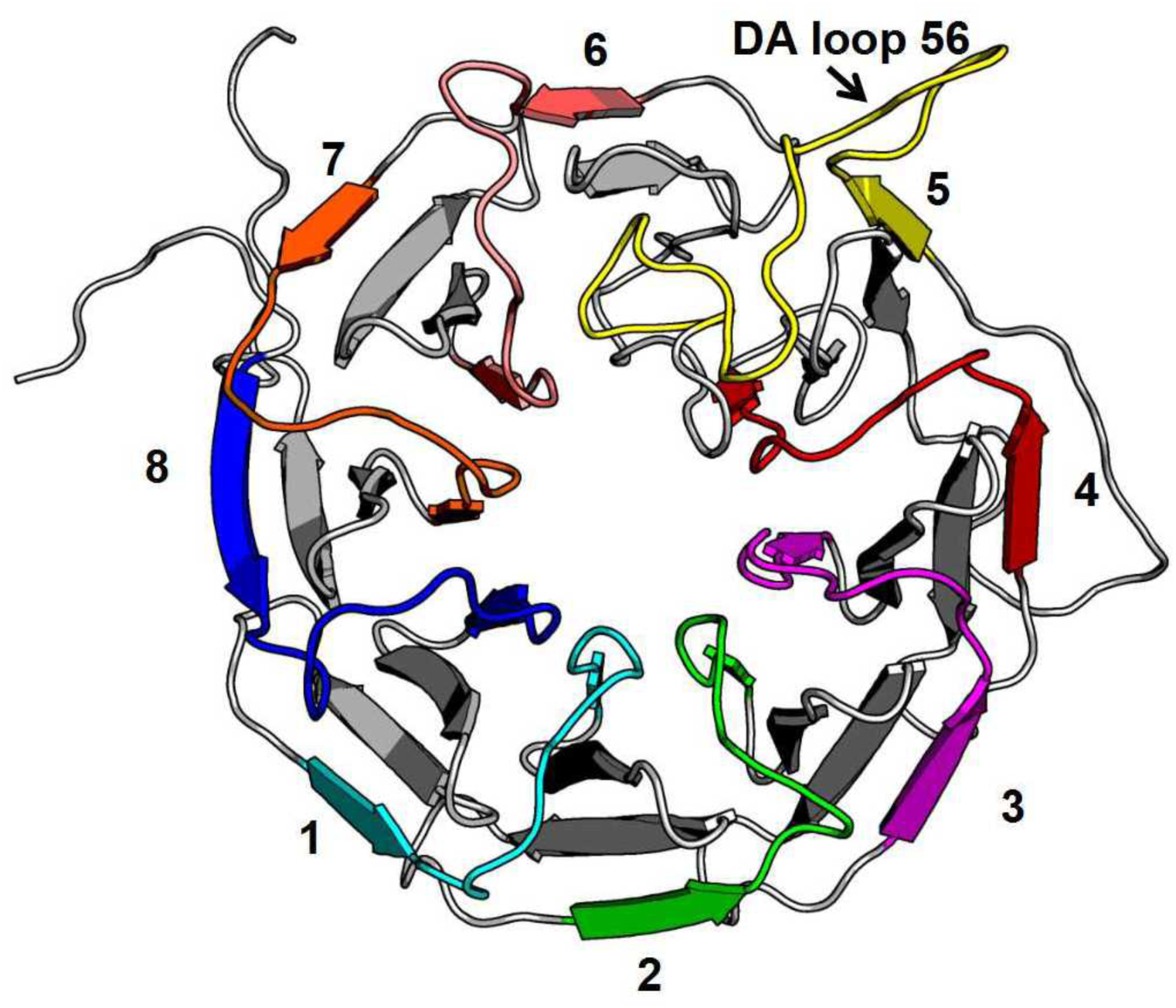
β-propeller blades constitute anti-parallel four β-sheets. Each β-propeller blades linked together with DA loop shown in different colors. DA loop connecting b 5 and 6 is longer and shown in yellow.

**Figure S3:**
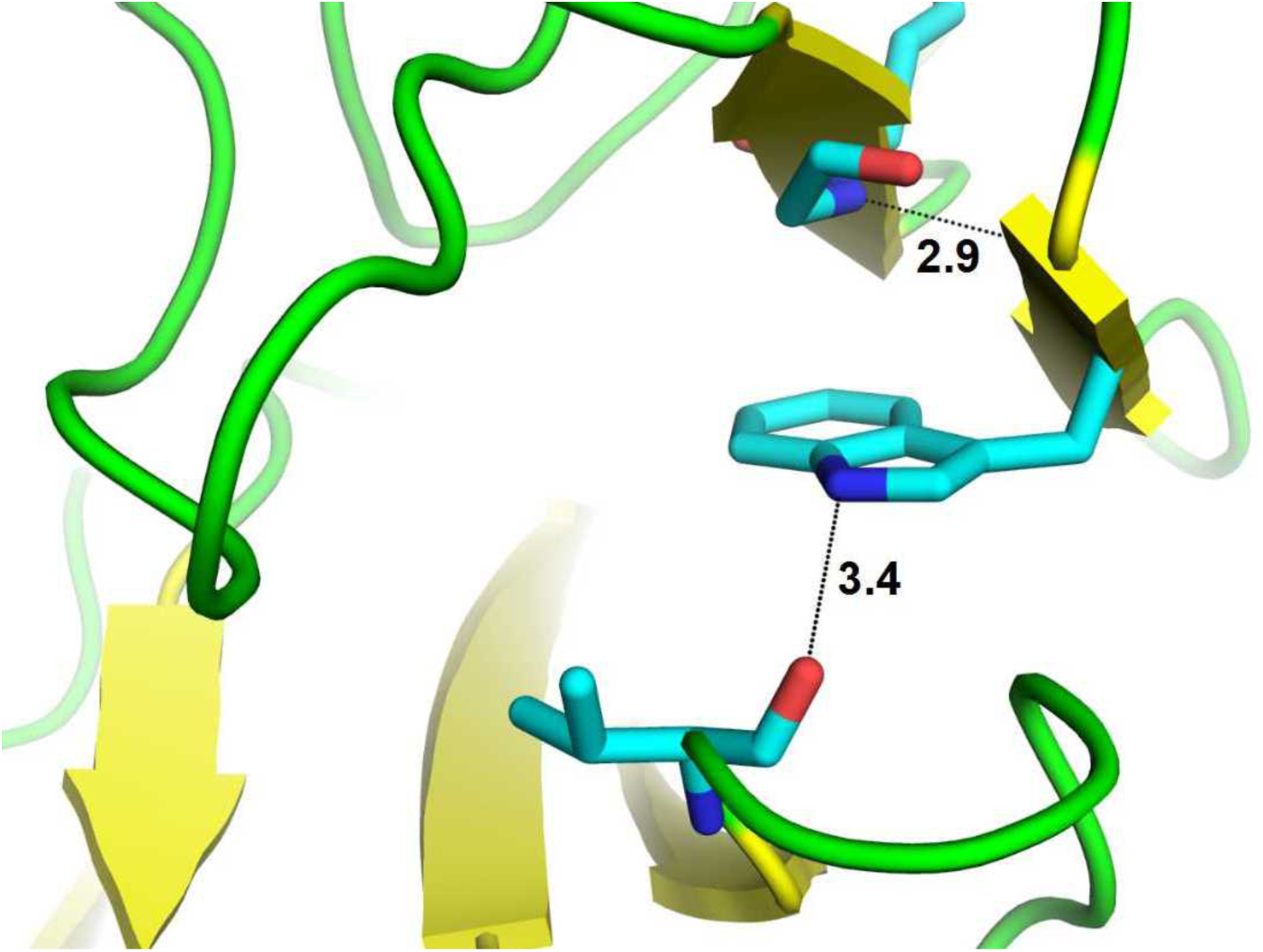
Hydrogen bonding interactions of conserved tryptophan in the D-strand with neighboring β-strands.

## References

1. Albrecht R, Zeth K. 2011. Structural basis of outer membrane protein biogenesis in bacteria. J Biol Chem 286:27792–803.

2. Akagawa M, Minematsu K, Shibata T, Kondo T, Ishii T, Uchida K. 2016. Identification of lactate dehydrogenase as a mammalian pyrroloquinoline quinone (PQQ)-binding protein. Sci Rep 6:26723.

3. Anthony C, Ghosh M.1998. The structure and function of the PQQ-containing quinoprotein dehydrogenases. Prog Biophys Mol Biol 69:1–21.

4. Anthony C, Ghosh M, Blake CC.1994. The structure and function of methanol dehydrogenase and related quinoproteins containing pyrrolo-quinoline quinone. Biochem J 304 (Pt 3):665–74.

5. Ashkenazy H, Abadi S, Martz E, Chay O, Mayrose I, Pupko T, Ben-Tal N. 2016. ConSurf 2016: an improved methodology to estimate and visualize evolutionary conservation in macromolecules. Nuc Aci Res W344–W350.

6. Barthe P, Mukamolova GV, Roumestand C, Cohen-Gonsaud M. 2010. The structure of PknB extracellular PASTA domain from mycobacterium tuberculosis suggests a ligand-dependent kinase activation. Structure 18:606–15.

7. Benkert P, Biasini M, Schwede T. 2011. Toward the estimation of the absolute quality of individual protein structure models. Bioinformatics 27: 343–350.

8. Bihani SC, Panicker L, Rajpurohit YS, Misra HS, Kumar V. 2018. drFrnE Represents a Hitherto Unknown Class of Eubacterial Cytoplasmic Disulfide Oxido-Reductases. Antioxid Redox Signal 28(4):296–310.

9. Blasius M, Sommer S, Hubscher U.2008. Deinococcus radiodurans: what belongs to the survival kit? Crit Rev Biochem Mol Biol 43:221–38.

10. Bolsunovsky A, Frolova T, Dementyev D, Sinitsyna O.2016. Low doses of gamma-radiation induce SOS response and increase mutation frequency in Escherichia coli and Salmonella typhimurium cells. Ecotoxicol Environ Saf 134P1:233–238.

11. Bonacossa de Almeida C, Coste G, Sommer S, Bailone A.2002. Quantification of RecA protein in Deinococcus radiodurans reveals involvement of RecA, but not LexA, in its regulation. Mol Genet Genomics 268:28–41.

12. Bourret RB, Silversmith RE.2010. Two-component signal transduction. Curr Opin Microbiol 13:113-5.

12. Chan, P.M., Ilangumaran, S., La Rose, J., Chakrabartty, A., and Rottapel, R. (2003) Autoinhibition of the kit receptor tyrosine kinase by the cytosolic juxtamembrane region. Mol Cell Biol 23: 3067–3078.

13. Cox MM, Battista JR.2005. Deinococcus radiodurans - the consummate survivor. Nat Rev Microbiol 3:882–92.

14. Daly MJ, Gaidamakova EK, Matrosova VY, Kiang JG, Fukumoto R, Lee DY, Wehr NB, Viteri GA, Berlett BS, Levine RL.2010. Small-molecule antioxidant proteome-shields in Deinococcus radiodurans. PLoS One 5:e12570.

15. Daly MJ, Minton KW.1996. An alternative pathway of recombination of chromosomal fragments precedes recA-dependent recombination in the radioresistant bacterium Deinococcus radiodurans. J Bacteriol 178:4461–71.

16. Desai, S.S., Rajpurohit, Y.S., Misra, H.S., and Deobagkar, D.N. (2011) Characterization of the role of the RadS/RadR two-component system in the radiation resistance of Deinococcus radiodurans. Microbiology 157: 2974–2982.

17. Didier JP, Cozzone AJ, Duclos B.2010. Phosphorylation of the virulence regulator SarA modulates its ability to bind DNA in Staphylococcus aureus. FEMS Microbiol Lett 306:30–6.

18. Dutta R, Qin L, Inouye M.1999. Histidine kinases: diversity of domain organization. Mol Microbiol 34:633–40.

19. Earl AM, Mohundro MM, Mian IS, Battista JR.2002. The IrrE protein of Deinococcus radiodurans R1 is a novel regulator of recA expression. J Bacteriol 184:6216–24.

20. Fernandez P, Saint-Joanis B, Barilone N, Jackson M, Gicquel B, Cole ST, Alzari PM.2006. The Ser/Thr protein kinase PknB is essential for sustaining mycobacterial growth. J Bacteriol 188:7778–84.

21. Gao R, Stock AM.2009. Biological insights from structures of two-component proteins. Annu Rev Microbiol 63:133–54.

22. Garcia-Garcia T, Poncet S, Derouiche A, Shi L, Mijakovic I, Noirot-Gros MF.2016. Role of Protein Phosphorylation in the Regulation of Cell Cycle and DNA-Related Processes in Bacteria. Front Microbiol 7:184.

23. Good MC, Greenstein AE, Young TA, Ng HL, Alber T.2004. Sensor domain of the Mycobacterium tuberculosis receptor Ser/Thr protein kinase, PknD, forms a highly symmetric beta propeller. J Mol Biol 339:459–69.

24. Griffith J, Black J, Faerman C, Swenson L, Wynn M, Lu F, Lippke J, Saxena K.2004. The structural basis for autoinhibition of FLT3 by the juxtamembrane domain. Mol Cell 13:169–78.

25. Hanks SK, Hunter T.1995. Protein kinases 6. The eukaryotic protein kinase superfamily: kinase (catalytic) domain structure and classification. FASEB J 9:576–96.

26. Hanks SK, Quinn AM, Hunter T.1988. The protein kinase family: conserved features and deduced phylogeny of the catalytic domains. Science 241:42–52.

27. Hu XJ, Li T, Wang Y, Xiong Y, Wu XH, Zhang DL, Ye ZQ, Wu YD.2017. Prokaryotic and Highly-Repetitive WD40 Proteins: A Systematic Study. Sci Rep 7:10585.

28. Hua Y, Narumi I, Gao G, Tian B, Satoh K, Kitayama S, Shen B.2003. PprI: a general switch responsible for extreme radioresistance of Deinococcus radiodurans. Biochem Biophys Res Commun 306:354–60.

29. Huse M, Kuriyan J.2002. The conformational plasticity of protein kinases. Cell 109:275–82.

30. Joshi B, Schmid R, Altendorf K, Apte SK.2004. Protein recycling is a major component of post-irradiation recovery in Deinococcus radiodurans strain R1. Biochem Biophys Res Commun 320:1112–7.

31. Källberg M,Wang H, Wang S, Peng J, Wang Z, Lu H, Xu J. 2012. Template-based protein structure modeling using the RaptorX web server. Nat Protocol 7, 1511–1522.

32. Kennelly PJ.2002. Protein kinases and protein phosphatases in prokaryotes: a genomic perspective. FEMS Microbiol Lett 206:1–8.

33. Khairnar NP, Misra HS, Apte SK.2003. Pyrroloquinoline-quinone synthesized in Escherichia coli by pyrroloquinoline-quinone synthase of Deinococcus radiodurans plays a role beyond mineral phosphate solubilization. Biochem Biophys Res Commun 312:303–8.

34. Kim JI, Cox MM.2002. The RecA proteins of Deinococcus radiodurans and Escherichia coli promote DNA strand exchange via inverse pathways. Proc Natl Acad Sci U S A 99:7917–21.

35. Kobir A, Shi L, Boskovic A, Grangeasse C, Franjevic D, Mijakovic I.2011. Protein phosphorylation in bacterial signal transduction. Biochim Biophys Acta 1810:989–94.

36. Krupa A, Srinivasan N.2005. Diversity in domain architectures of Ser/Thr kinases and their homologues in prokaryotes. BMC Genomics 6:129.

37. Laskowski RA, MacArthur MW, Moss DS, Thornton, J. M. (1993) PROCHECK - a program to check the stereochemical quality of protein structures. J App Cry 26: 283–291.

38. Leonard, C.J., Aravind, L., and Koonin, E.V. 1998. Novel families of putative protein kinases in bacteria and archaea: evolution of the “eukaryotic” protein kinase superfamily. Genome Res 8: 1038–1047.

39. Liu Y, Zhou J, Omelchenko MV, Beliaev AS, Venkateswaran A, Stair J, Wu L, Thompson DK, Xu D, Rogozin IB, Gaidamakova EK, Zhai M, Makarova KS, Koonin EV, Daly MJ.2003. Transcriptome dynamics of Deinococcus radiodurans recovering from ionizing radiation. Proc Natl Acad Sci U S A 100:4191–6.

40. Macek B, Mijakovic I, Olsen JV, Gnad F, Kumar C, Jensen PR, Mann M.2007. The serine/threonine/tyrosine phosphoproteome of the model bacterium Bacillus subtilis. Mol Cell Proteomics 6:697–707.

41. Maurya GK, Modi K, Misra HS. 2016. Divisome and segrosome components of Deinococcus radiodurans interact through cell division regulatory proteins. Microbiol 162(8):1321–1334.

42. Maurya GK, Modi K, Banerjee M, Chaudhary R, Rajpurohit YS, Misra HS.2018. Phosphorylation of FtsZ and FtsA by a DNA Damage-Responsive Ser/Thr Protein Kinase Affects Their Functional Interactions in Deinococcus radiodurans. mSphere 3.

43. Minton KW.1994. DNA repair in the extremely radioresistant bacterium Deinococcus radiodurans. Mol Microbiol 13:9–15.

44. Misra HS, Khairnar NP, Barik A, Indira Priyadarsini K, Mohan H, Apte SK.2004. Pyrroloquinoline-quinone: a reactive oxygen species scavenger in bacteria. FEBS Lett 578:26–30.

45. Misra HS, Rajpurohit YS, Khairnar NP. 2012. Pyrroloquinoline-quinone and its versatile roles in biological processes. J Basic Microbiol 53(6):518–31.

46. Misra H.S., Rajpurohit, Y.S., and Kota, S. (2013) Physiological and molecular basis of extreme radioresistance in Deinococcus radiodurans Current Sci 104:194–205.

47. Misra HS, Maurya GK, Chaudhary R, Misra CS. 2018. Interdependence of bacterial cell division and genome segregation and its potential in drug development. Microbiol Res 208:12–24.

48. Mishra S, Chaudhary R, Singh S, Kota S, Misra HS. 2019. Guanine Quadruplex DNA Regulates Gamma Radiation Response of Genome Functions in the Radioresistant Bacterium *Deinococcus radiodurans*. J Bacteriol 8: 201(17).

49. Molle V, Kremer L.2010. Division and cell envelope regulation by Ser/Thr phosphorylation: Mycobacterium shows the way. Mol Microbiol 75:1064–77.

50. Mylona A, Fernandez-Tornero C, Legrand P, Haupt M, Sentenac A, Acker J, Muller CW.2006. Structure of the tau60/Delta tau91 subcomplex of yeast transcription factor IIIC: insights into preinitiation complex assembly. Mol Cell 24:221–32.

51. Narumi I, Satoh K, Kikuchi M, Funayama T, Yanagisawa T, Kobayashi Y, Watanabe H, Yamamoto K.2001. The LexA protein from Deinococcus radiodurans is not involved in RecA induction following gamma irradiation. J Bacteriol 183:6951–6.

52. Oshima T, Aiba H, Masuda Y, Kanaya S, Sugiura M, Wanner BL, Mori H, Mizuno T.2002. Transcriptome analysis of all two-component regulatory system mutants of Escherichia coli K-12. Mol Microbiol 46:281–91.

53. Parkinson JS.1993. Signal transduction schemes of bacteria. Cell 73:857–71.

54. Pereira SF, Goss L, Dworkin J.2011. Eukaryote-like serine/threonine kinases and phosphatases in bacteria. Microbiol Mol Biol Rev 75:192–212.

55. Rajpurohit YS, Bihani SC, Waldor MK, Misra HS.2016. Phosphorylation of Deinococcus radiodurans RecA Regulates Its Activity and May Contribute to Radioresistance. J Biol Chem 291:16672–85.

56. Rajpurohit YS, Gopalakrishnan R, Misra HS.2008. Involvement of a protein kinase activity inducer in DNA double strand break repair and radioresistance of Deinococcus radiodurans. J Bacteriol 190:3948–54.

57. Rajpurohit YS, Misra HS.2010. Characterization of a DNA damage-inducible membrane protein kinase from Deinococcus radiodurans and its role in bacterial radioresistance and DNA strand break repair. Mol Microbiol 77:1470–82.

58. Rajpurohit YS, Misra HS.2013. Structure-function study of deinococcal serine/threonine protein kinase implicates its kinase activity and DNA repair protein phosphorylation roles in radioresistance of Deinococcus radiodurans. Int J Biochem Cell Biol 45:2541–52.

59. Rajpurohit YS, Misra HS. 2013b. DR1769, a protein with N-terminal beta propeller repeats and a low-complexity hydrophilic tail, plays a role in desiccation tolerance of Deinococcus radiodurans. J Bacteriol 195 (17):3888-3896.

60. Rajpurohit YS, Desai SS, Misra HS. 2013c. Pyrroloquinoline quinone and a quinoprotein kinase support γ-radiation resistance in Deinococcus radiodurans and regulate gene expression. J Basic Microbiol 53(6):518–31.

61. Reisz JA, Bansal N, Qian J, Zhao W, Furdui CM. 2014. Effects of ionizing radiation on biological molecules--mechanisms of damage and emerging methods of detection. Anti Red Sig 21(2):260–92.

62. Sancar A, Lindsey-Boltz LA, Unsal-Kacmaz K, Linn S.2004. Molecular mechanisms of mammalian DNA repair and the DNA damage checkpoints. Annu Rev Biochem 73:39–85.

63. Sassetti CM, Boyd DH, Rubin EJ.2003. Genes required for mycobacterial growth defined by high density mutagenesis. Mol Microbiol 48:77–84.

64. Scherr N, Honnappa S, Kunz G, Mueller P, Jayachandran R, Winkler F, Pieters J, Steinmetz MO.2007. Structural basis for the specific inhibition of protein kinase G, a virulence factor of Mycobacterium tuberculosis. Proc Natl Acad Sci U S A 104:12151–6.

65. Schrover JM, Frank J, van Wielink JE, Duine JA.1993. Quaternary structure of quinoprotein ethanol dehydrogenase from Pseudomonas aeruginosa and its reoxidation with a novel cytochrome c from this organism. Biochem J 290 (Pt 1):123–7.

66. Shah IM, Laaberki MH, Popham DL, Dworkin J.2008. A eukaryotic-like Ser/Thr kinase signals bacteria to exit dormancy in response to peptidoglycan fragments. Cell 135:486–96.

67. Sharma DK, Siddiqui MQ, Gadewal N, Choudhary RK, Varma AK, Misra HS, Rajpurohit YS.2020. Phosphorylation of deinococcal RecA affects its structural and functional dynamics implicated for its roles in radioresistance of Deinococcus radiodurans. J Biomol Struct Dyn 38:114–123.

68. Shimoni Y, Altuvia S, Margalit H, Biham O.2009. Stochastic analysis of the SOS response in Escherichia coli. PLoS One 4:e5363.

69. Shuid AN, Kempster R, McGuffin LJ. 2017. ReFOLD: a server for the refinement of 3D protein models guided by accurate quality estimates. Nuc Aci Res 3: 45.

70. Skaar JR, Pagan JK, Pagano M.2014. SCF ubiquitin ligase-targeted therapies. Nat Rev Drug Discov 13:889–903.

71. Slade D, Lindner AB, Paul G, Radman M.2009. Recombination and replication in DNA repair of heavily irradiated Deinococcus radiodurans. Cell 136:1044–55.

72. Smith TA, Kirkpatrick DR, Smith S, Smith TK, et al. 2017. Radioprotective agents to prevent cellular damage due to ionizing radiation. J Trans Med 15: 232.

73. Tanaka M, Earl AM, Howell HA, Park MJ, Eisen JA, Peterson SN, Battista JR.2004. Analysis of Deinococcus radiodurans’s transcriptional response to ionizing radiation and desiccation reveals novel proteins that contribute to extreme radioresistance. Genetics 168:21–33.

74. Wall MA, Coleman DE, Lee E, Iniguez-Lluhi JA, Posner BA, Gilman AG, Sprang SR.1995. The structure of the G protein heterotrimer Gi alpha 1 beta 1 gamma 2. Cell 83:1047–58.

75. Wang L, Xu G, Chen H, Zhao Y, Xu N, Tian B, Hua Y. 2008. DrRRA: a novel response regulator essential for the extreme radioresistance of Deinococcus radiodurans. Mol Microbiol 67: 1211–1222.

76. Wiederstein M, Sippl MJ. 2007. ProSA-web: interactive web service for the recognition of errors in three-dimensional structures of proteins. Nuc Aci Res 35: W407–410.

77. Wu XH, Wang Y, Zhuo Z, Jiang F, Wu YD.2012. Identifying the hotspots on the top faces of WD40-repeat proteins from their primary sequences by beta-bulges and DHSW tetrads. PLoS One 7:e43005.

78. Wybenga-Groot LE, Baskin B, Ong SH, Tong J, Pawson T, Sicheri F.2001. Structural basis for autoinhibition of the Ephb2 receptor tyrosine kinase by the unphosphorylated juxtamembrane region. Cell 106:745–57.

79. Zahradka K, Slade D, Bailone A, Sommer S, Averbeck D, Petranovic M, Lindner AB, Radman M.2006. Reassembly of shattered chromosomes in Deinococcus radiodurans. Nature 443:569–73.

80. Zhang C, Zhang F.2015. The Multifunctions of WD40 Proteins in Genome Integrity and Cell Cycle Progression. J Genomics 3:40–50.

81. Zhou BB, Elledge SJ.2000. The DNA damage response: putting checkpoints in perspective. Nature 408:433–9.

